# (*S*)-ketamine augments behavioral and physiological responses to Δ^9^-tetrahydrocannabinol

**DOI:** 10.64898/2025.12.18.695060

**Authors:** Erik Keimpema, Valerie Vigl, Natalya Torgasheva, Roman A. Romanov, Siegfried Kasper, Tibor Harkany

## Abstract

(*S*)-ketamine is a fast-acting therapeutic to manage treatment-resistant depression as well as psychiatric emergencies. As the non-prescribed use of ketamine has rapidly increased over the past decade, its possible interactions with other recreational drugs are of increasing interest. Here, we addressed if Δ^9^-tetrahydrocannabinol (THC), the main psychoactive constituent of *Cannabis sp*., and (*S*)-ketamine could affect each other’s psychoactive profile in a pharmacological triad. In healthy male mice, the sequential use of (*S*)-ketamine followed by THC at human-relevant doses did not result in either synergistic or antagonistic effects acutely. In contrast, THC-induced hypothermia was nearly doubled by subsequent (*S*)-ketamine administration. THC-induced hypolocomotion was not only prolonged by (*S*)-ketamine, but axial rotation and stumbling during ambulation in an open field were increased in subjects exposed sequentially to both drugs. Likewise, the performance of subjects exposed to THC and then (*S*)-ketamine worsened on a rotarod. Combined drug treatment led to unusual epinephrine and corticosterone profiles in blood plasma. We suggest that (*S*)-ketamine augments adverse THC effects, which at the endocrine level invoke the acute deregulation of the hypothalamus-pituitary-adrenal axis. Our findings support the clinical notion that combining cannabis and (*S*)-ketamine should be avoided.

## Introduction

In recent years, ketamine gained traction as a safe therapeutic option for patients with treatment-resistant depression (TRD)^1–4^. Unlike conventional antidepressants, including slow-acting tetracyclics and selective serotonin/epinephrine reuptake inhibitors (SSRIs/SNRIs), both racemic ketamine and its more powerful (*S*)-enantiomer exhibit rapid and robust antidepressant effects, which are sustained for up to weeks post-treatment^5–7^. Its fast action is due to antagonism at N-methyl-D-aspartate receptors (NMDARs), which is valuable to treat psychiatric emergencies and to instantaneously offset self-harm^8^. Accordingly, the clinical use of (*S*)-ketamine has been approved by health care systems around the world, both as intravenous and over-the-counter intranasal formulations^1,9^. Despite its therapeutic efficacy and perceived safety, (*S*)-ketamine can cause acute negative side effects including dizziness, balance and movement dyscoordination, dissociation, increased blood pressure, and nausea^7^. Chronic ketamine use is paradoxically associated with the development of anxiety, depression, cognitive defects, and persisting psychosis^10^. As such, its long-term therapeutic benefits and safety profile are still questioned both in clinical settings^11^ and upon recreational use^12^, especially considering its low-to-moderate addictive qualities^13^. This complexity of action is also notable because of (*S*)-ketamine’s interaction with other pharmaceutical agents. For instance, benzodiazepines, mood stabilizers, and anti-psychotics, alternative treatment options for anxiety and depression^14^, interfere with the antidepressant effects of ketamine therapy^15^. Even more concerning is that the non-prescribed use of ketamine has significantly risen in the past decade because of both recreational use and personal self-medication^16,17^. Thus, a significant gap of knowledge exists on how recreational drugs interact with (*S*)-ketamine^18^, and if adverse outcomes might prevail.

A dominant candidate for recreational interactions is cannabis because its use has similarly increased over the past decade due to legislative changes^19^ along with plant selection for increased psychoactive action^20^. Additionally, cannabis is often used to self-medicate anxiety^21^ and depression^1,22^, and to manage pain^23^. Nevertheless, excessive cannabis use can precipitate mental illnesses, such as schizophrenia and psychosis^24^. Considering that Δ^9^-tetrahydrocannabinol (THC), the main psychoactive constituent in *Cannabis sp*., acts at metabotropic G protein-coupled cannabinoid receptors to induce protracted intracellular signaling events^25,26^, its temporal window of efficacy is expected to be beyond that of (*S*)-ketamine in the nervous system, at least acutely.

Here, we addressed if (*S*)-ketamine could acutely interact with THC at doses for both drugs relevant to human recreational exposure. By using a physiological triad encompassing the monitoring of core body temperature, locomotion, and motor coordination in healthy mice^27^, we first showed that (*S*)-ketamine augmented THC-induced hypothermia. Next, (*S*)-ketamine was not only found to lose its anxiolytic effect in an open field arena^28–30^ when administered to THC-exposed subjects, but also dose-dependently prolong THC-induced hypolocomotion. Similarly, subjects exposed to THC-then-(*S*)-ketamine doses had significantly worsened motor coordination, as compared to the outcome of single drug use. No interaction was observed when reversing the sequence of (*S*)-ketamine and THC in any test. We symptomatically attributed the worsening of behavioral outcomes, at least in part, to THC preventing the (*S*)-ketamine-induced reduction of circulating adrenocorticotropin-releasing hormone, epinephrine, and corticosterone levels required for effective anxiolysis^31,32^. Single-cell RNA-seq of the HPA axis suggested drug interactions to dominate at the level of corticotropin-releasing hormone (CRH)-containing neurons in the paraventricular hypothalamus. In sum, we suggest that THC and (*S*)-ketamine interact at behavioral and endocrine levels. Thus, our data support that cannabis use before (*S*)-ketamine sessions should be avoided, especially noting the long half-life of THC metabolites in the human body^26^.

## Results

### Validation of (*S*)-ketamine action in male mice

As drug effects differ with animal strain, sex, and age^33–35^, we first constructed a dose-response relationship for (*S*)-ketamine in cohorts of male C57BL/6JRj mice (*experimental design:* **Fig. S1a**). For (*S*)-ketamine (Eskelan, GL Pharma), doses of 5, 10, and 15 mg/kg were tested, which translated to 28–85 mg for an average adult human of 70 kg^36,37^, thus encompassing the 0.5 mg/kg concentration preferred for the clinical treatment of depression^38^.

Firstly, we observed a dose-dependent modification of core body temperature 30 min after (*S*)- ketamine injection (**Fig. S2a**)^39^, with 10 mg/kg (*S*)-ketamine evoking a statistically significant temperature drop (0.67 ± 0.19 °C [vehicle; *n* = 10] *vs.* -0.60 ± 0.31 °C [(*S*)-ketamine; *n* = 4]; *p* < 0.001). Secondly, (*S*)-ketamine did not affect locomotion in an open field over 30 min (**Fig. S2b,c**). Nevertheless, when binning ambulatory activity per 5 min intervals, mice displayed hypermobility during the initial 5-min epoch immediately after exposure to 10 mg/kg (*S*)-ketamine, as compared to controls (548 ± 21 cm [vehicle; *n* = 10] *vs.* 657 ± 33 cm [(*S*)-ketamine; *n* = 5]; *p* < 0.001; **Fig. S2b_1_**). Fifteen mg/kg (*S*)-ketamine had reduced efficacy, which we attributed to its slowing of motor movements^28^. We also observed stereotypic axial rotations^40,41^ at (*S*)-ketamine concentrations ≥10 mg/kg (0.04 ± 0.02 rotations/m [vehicle; *n* = 10] *vs.* 0.40 ± 0.11 rotations/m [(*S*)-ketamine 10 mg/kg; *n* = 5]; *p* < 0.008; **Fig. S2d**). Grooming was not affected (**Fig. S2e**). In contrast, both supported and unsupported rearing were dose-dependently reduced at (*S*)-ketamine doses ≥ 10 mg/kg (*supported*: 15.7 ± 1.3% [vehicle; *n* = 10] *vs.* 4.2 ± 0.9% [(*S*)-ketamine 10 mg/kg; *n* = 5]; *p* < 0.001; *unsupported*: 8.4 ± 0.7% [vehicle; *n* = 10] *vs.* 5.0 ± 1.0% [(*S*)-ketamine 10 mg/kg; *n* = 5]; data were expressed as percentage of total time; *p* < 0.001; **Fig. S2e,e_1_**).

Next, we replicated the above experiments after 60 min, in line with findings of a recent report^29^: core body temperature was normal (**Fig. S3a**), alike exploratory (**Fig. S3b-c**), and maintenance behaviors (**Fig. S3d-e_2_**). Initial hyperlocomotion (5 min bin) was no longer measured at any (*S*)- ketamine concentration. Instead, moderate hypolocomotion was present in mice treated with 15 mg/kg (*S*)-ketamine (517 ± 29 cm [vehicle; *n* = 10] *vs.* 342 ± 23 cm [(*S*)-ketamine; *n* = 5]; *p* < 0.001; **Fig. S3b_1_**), likely due to the drug’s sedative properties. Consequently, we deemed (*S*)- ketamine fast-acting, with its acute effects limited to a 30-min period, and selected its 10 mg/kg concentration for subsequent experiments.

### Validation of THC action in male mice

In a separate set of experiments, we injected 1, 3, 5 or 10 mg/kg synthetic THC (Dronabinol, Gatt-Koller), which translated to 5.7–57 mg for an average adult human weighing 70 kg^36,37^, encompassing typical THC concentrations consumed when using cannabis cigarettes and edibles^42,43^. As above, and in conformity with the ‘tetrad test’^27^, core body temperature, locomotor activity, and motor coordination were assessed 30 min after drug delivery (**Fig. S1b**).

Firstly, we found a biologically-relevant decrease in core body temperature at THC doses of ≥5 mg/kg, as compared to vehicle (0.16 ± 0.32 °C [vehicle; *n* = 9] *vs.* -2.0 ± 0.29 °C [5 mg/kg THC; *n* = 7]; *p* = 0.003; **Fig. S4a**)^27^. Secondly, THC induced dose-dependent hypolocomotion in the open field (128 ± 6 m [vehicle; *n* = 10] *vs.* 94 ± 5 m [1 mg/kg THC; *n* = 3]; *p* = 0.04; 77 ± 3 m [3 mg/kg THC; *n* = 3]; *p* = 0.007; 28 ± 6 m [5 mg/kg THC; *n* = 7]; *p* < 0.001; 16 ± 8 m [10 mg/kg THC; *n* = 4]; *p* < 0.001; **Fig. S4b,c**). When binned for 5-min intervals, hypolocomotion persisted throughout the 30-min period (**Fig. S4b_1_**). Axial rotations were not observed (**Fig. S4d**). Grooming (**Fig. S4e**) was markedly reduced at THC doses of >3 mg/kg (**Fig. S4e**), while both supported and unsupported rearing were significantly reduced in a dose-dependent fashion (**Fig. S4e_1_,e_2_**).

In the catalepsy test, mice were generally classified as affected at THC concentrations of ≥ 5 mg/kg. However, the ‘pop-corn’ effect (a sudden intense startle response)^27^ at higher THC concentrations made it difficult to quantify the time the mice spent on the bar because they jumped away immediately when placed onto it (**Video S1*-*S3**). Based on these findings, 5 mg/kg THC was taken as the concentration of choice in follow-up experiments.

### Interaction when (*S*)-ketamine preceded THC

We tested if pre-treatment (30 min) with 10 mg/kg (*S*)-ketamine could modulate the behavioral effects of 5 mg/kg THC on core body temperature, locomotion, and motor co-ordination (*experimental design:* **Fig. S1c**). For any of the concentrations tested, (*S*)-ketamine did not affect THC-induced changes in core body temperature (-2.44 ± 0.58 °C [THC] *vs.* -3.40 ± 0.73 °C [5 mg/kg (*S*)-ketamine + THC]; *p* = 0.58; -2.40 ± 0.46 °C [10 mg/kg (*S*)-ketamine + THC]; *p* = 0.5 and -3.15 ± 0.73 °C [15 mg/kg (*S*)-ketamine + THC]; *p* = 0.98; **Fig. 1a**), locomotion (17.06 ± 4.11 m [THC] *vs.* 9.66 ± 1.89 m [5 mg/kg (*S*)-ketamine + THC]; *p* = 0.63; 12.61 ± 3.25 m [10 mg/kg (*S*)-ketamine + THC]; *p* = 0.77 and 10.71 ± 2.81 m [15 mg/kg (*S*)-ketamine + THC]; *p* = 0.63; **Fig. 1b-c**, including 5-min bins), maintenance (1.85 ± 0.63% [THC] *vs.* 2.47 ± 0.88% [5 mg/kg (*S*)-ketamine + THC]; *p* = 0.73; 1.82 ± 0.59% [10 mg/kg (*S*)-ketamine + THC]; *p* = 0.99; 1.99 ± 0.11% [15 mg/kg (*S*)-ketamine + THC]; *p* = 0.94; data were expressed as percentage of total time; **Fig. 1e**), and exploratory behaviors (**Fig. 1e_1_,e_2_**). Axial rotations otherwise seen for 10 mg/kg (*S*)-ketamine alone (**Fig. S2d**) were abolished by the THC dose tested (0 ± 0 rotations/m [THC] *vs.* 0.0 ± 0.0 rotations/m [5 mg/kg (*S*)-ketamine + THC]; *p* = 0.99; 0.01 ± 0.01 rotations/m [10 mg/kg (*S*)-ketamine + THC]; *p* = 0.57, and 0.03 ± 0.03 rotations/m [15 mg/kg (*S*)-ketamine + THC]; *p* = 0.11; **Fig. 1d**). Thus, we concluded that pre-treatment with (*S*)-ketamine does not significantly impact subsequent THC effects, presumable due to its short half-life^44^.

**Figure 1.**
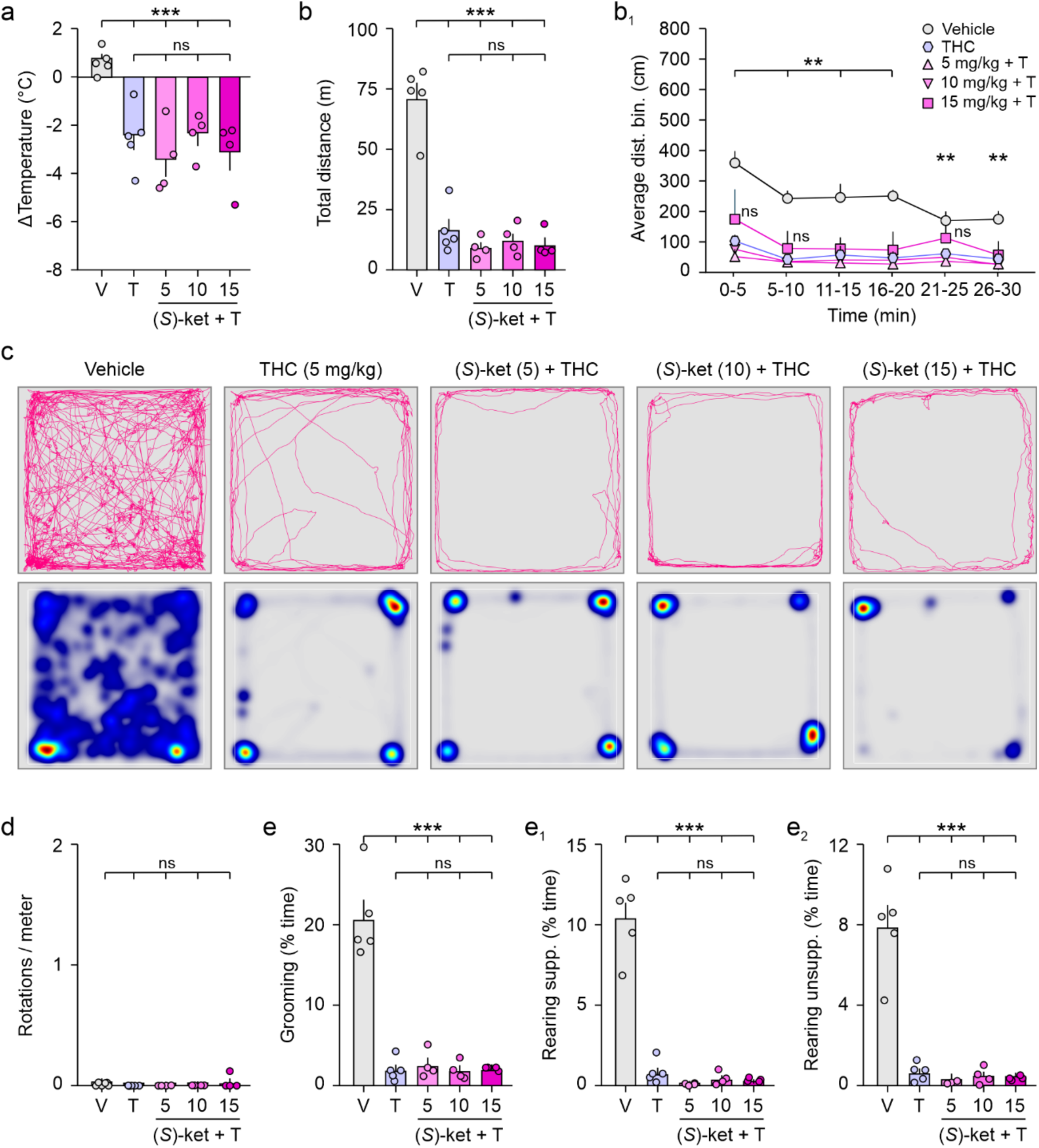
(*S*)-ketamine pretreatment did not modify THC effects. (**a**) Changes in core body temperature 60 min after (*S*)-ketamine, that is, 30 min after THC exposure (*n* = 4-5 / group). (**b,b_1_**) Distance travelled in an open field test for 30 min. (*S*)-ketamine pretreatment did not modify THC-induced hypolocomotion. (**c**) Representative trajectories (*top*) and heat maps of time spent at specific locations in an open field (*bottom*; blue-to-red color scale refers to locations with none-to-maximum occupancy). (**d**) Number of axial rotations (‘circling behavior’). (**e-e_2_**) Grooming and rearing behavior during a 30-min period in the open field. Colored circles denote individual data points. Male mice were used throughout. Bar graphs show means ± s.e.m. \**p* < 0.05; \*\**p* < 0.01; \*\*\**p* < 0.001; ns, non-significant. *Abbreviations*: (*S*)-ket, (*S*)-ketamine; (*S*)-ket + T, (*S*)- ketamine + THC at the concentrations shown (in mg/kg); T, THC; V, vehicle.

### Interaction when THC preceded (*S*)-ketamine

Next, we reversed the order in which the drugs were administered (*experimental design*: **Fig. S1d**). When THC (5 mg/kg) preceded any of the (*S*)-ketamine doses by 30 min, the change in core body temperature nearly doubled (-2.41 ± 0.41 °C [THC] *vs.* -5.15 ± 0.28 °C [THC + 5 mg/kg (*S*)-ketamine], *p* < 0.001; -5.15 ± 0.61 °C [THC + 10 mg/kg (*S*)-ketamine], *p* < 0.001; -5.20 ± 0.33°C [THC + 15 mg/kg (*S*)-ketamine], *p* < 0.001; *n* = 4-6/group; **Fig. 2a**). We suggest that the combination of 5 mg THC + 5 mg/kg (*S*)- ketamine already reached physiologically tolerable maximum, with higher doses showing a ‘ceiling effect’ (see **Fig. S5a-a_2_** for a comparison between experiments).

**Figure 2.**
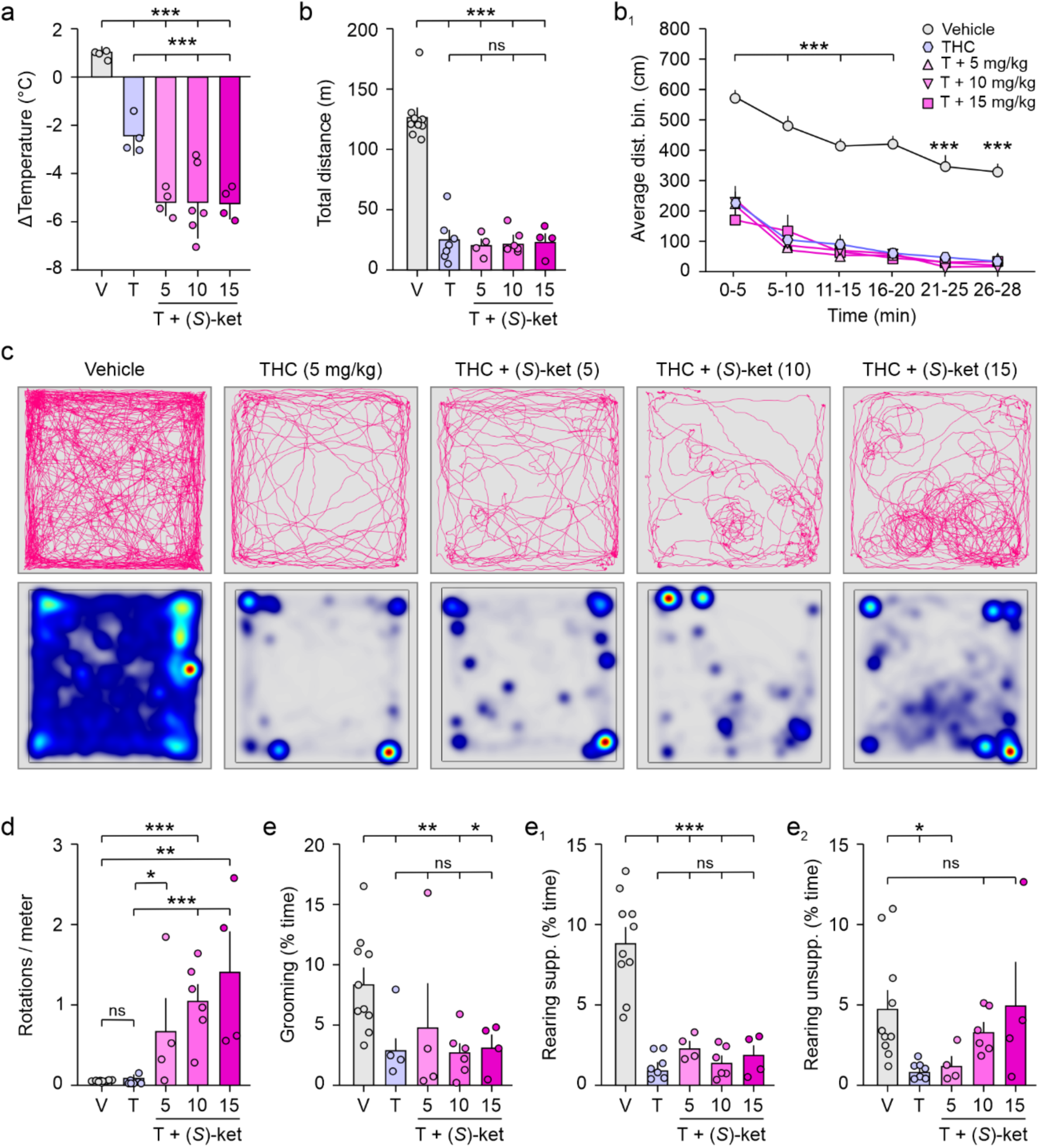
Interaction between THC followed by (*S*)-ketamine. (**a**) Differences in core body temperature 60 min after THC exposure; that is 30 min after (*S*)-ketamine (*n* = 4-6/group). (**b,b_1_**) Distance travelled in an open field test immediately after (*S*)-ketamine infusion. (**c**) Representative trajectories (*top*) and heat maps of time spent at specific locations in an open field (*bottom*; blue-to-red color scale refers to locations with none-to-maximum occupancy). (**d**) Number of axial rotations (‘circling behavior’), which increased as a factor of the (*S*)-ketamine concentrations (*n* = 4-10/group). (**e-e_2_**) Grooming and rearing behavior during a 30-min period in the open field. Colored circles denote individual data points in male mice. Bar graphs show means ± s.e.m. \**p* < 0.05; \*\**p* < 0.01; \*\*\**p* < 0.001; ns, non-significant. *Abbreviations*: (*S*)-ket, (*S*)-ketamine; T + (*S*)-ket, THC + (*S*)-ketamine at the concentrations shown (in mg/kg); T, THC; V, vehicle.

The total distance of ambulation in an open field was not modified by any of the (*S*)-ketamine doses in mice pretreated with 5 mg/kg THC. Notably, the initial hypermobility (0-5 min bin) characteristic of fast-acting (*S*)-ketamine was no longer seen with THC pretreatment (577 ± 15 cm [vehicle] *vs*. 229 ± 53 cm [THC], *p* < 0. 001 *vs.* 219 ± 45 cm [THC + 10 mg/kg (*S*)-ketamine], *p* = 0.9 (*vs.* THC); *n* = 4-10; **Fig. 2b_1_ *vs.* Fig. S2b_1_; and Fig. S5b,b_1_** for comparison).

At the same time, axial rotations, which we interpreted as a marker of disrupted motor coordination, were augmented dose-dependently (0.03 ± 0.01 rotation/m [vehicle] *vs*. 0.04 ± 0.02 rotation/m [THC]; *p* = 0.99 *vs.* 0.68 ± 0.39 rotation/m [THC + 5 mg/kg (*S*)-ketamine]; *p* = 0.15; 1.05 ± 0.19 rotation/m [THC + 10 mg/kg (*S*)-ketamine]; *p* = 0.017, and 1.42 ± 0.5 rotation/m [THC + 15 mg/kg (*S*)-ketamine]; *p* = 0.005; *n* = 4-10/group; also see movement traces in **Fig. 2c,d** and **Fig. 5c-c_2_** for comparison), reminiscent of the effects of much higher (*S*)-ketamine doses (≥30 mg/kg)^40,41^. Neither grooming nor supported rearing were improved. Yet escalating (*S*)-ketamine doses partly improved unsupported rearing, as compared to THC alone (**Fig. 2e-e_2_**).

The balancing deficits (axis rotations) together with occasional tripping and stumbling (**Video S4**) when increasing the concentration of (*S*)-ketamine worsened upon THC pretreatment. This observation prompted us to test motor coordination on a rotarod. We found that neither THC nor (*S*)-ketamine groups performed worse than their respective (vehicle) controls 1 h post (*S*)-ketamine exposure (157 ± 18 sec [vehicle] *vs*. 152 ± 27 sec [THC]; *p* = 0.90, and 128 ± 25 sec [(*S*)-ketamine]; *p* = 0.40; *n* = 4-6/group; **Fig. 3a**),. Nevertheless, subjects exposed to combined drug treatment had significantly more falls and failures (157 ± 18 sec [vehicle] *vs*. 98 ± 17 sec [THC + 10 mg/kg (*S*)-ketamine]; *p* = 0.025; *n* = 4-8/group; **Fig. 3a**), reinforcing an adverse interaction between THC and (*S*)-ketamine.

**Figure 3.**
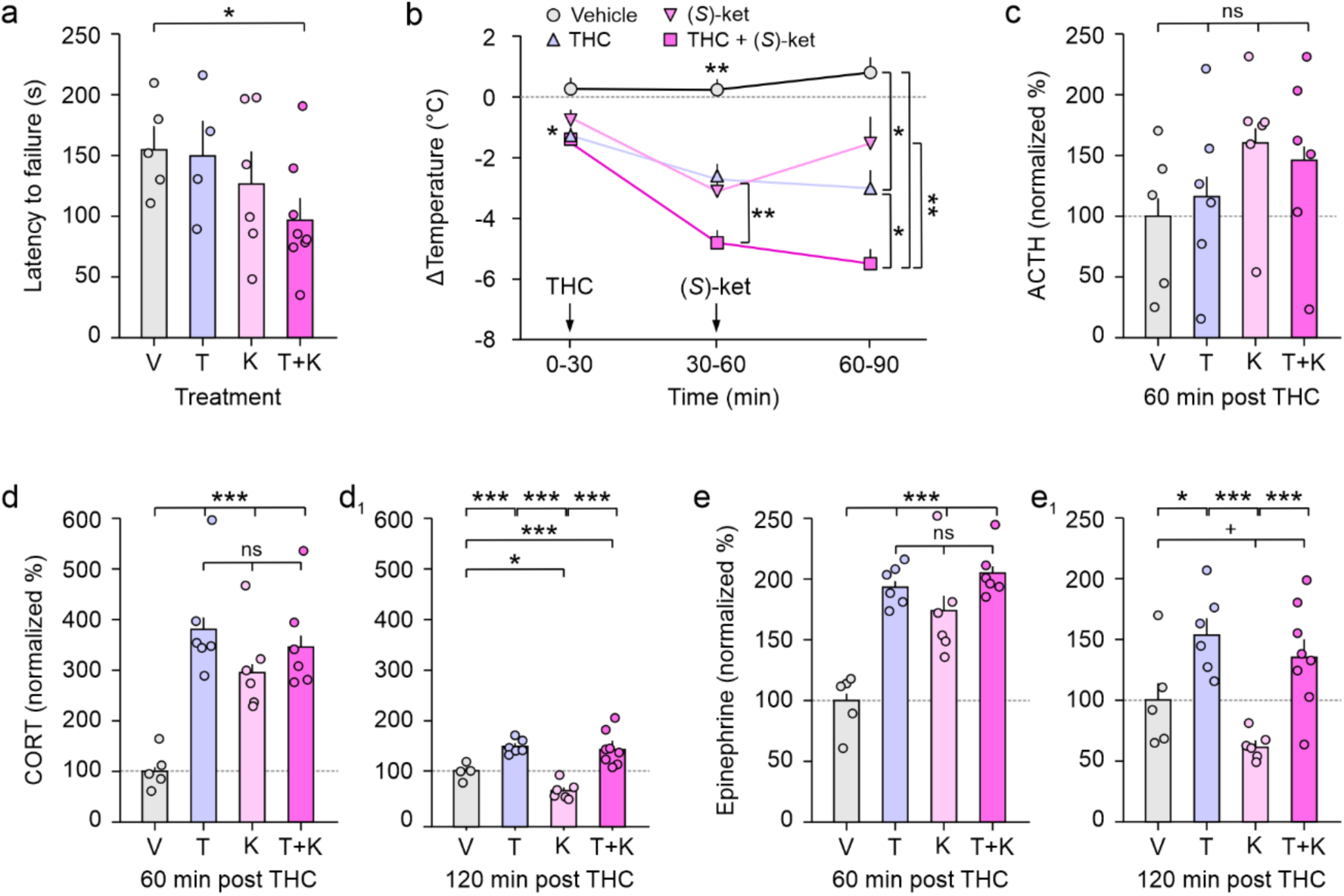
Behavioral, temperature, and serum hormone changes after combined treatment. (**a**) Latency to falling off (or holding on for a single rotation) of a rotarod device. (**b**) Changes in body temperature over time (*n* = 4-6/group). Note that core body temperature does not recover in the THC + (*S*)-ketamine group as compared to (*S*)-ketamine alone. (**c-e**) Plasma levels of adrenocorticotropic hormone (ACTH, *c*), corticosterone (CORT, *d*) and epinephrine (*e*) 60 or 120 min after THC injection (*n* = 5-8/group). (*S*)-ketamine was given 30 min after THC application. Data in male mice were normalized to the average of the vehicle group (horizontal dashed line). ^+^*p* < 0.1; \**p* < 0.05; \*\**p* < 0.01; \*\*\**p* < 0.001; ns, non-significant. *Abbreviations*: (*S*)-ket, (*S*)-ketamine; T + K, THC + (*S*)-ketamine at the concentrations shown (in mg/kg); T, THC; V, vehicle.

### Synergistic effects of THC and (*S*)-ketamine on core temperature

To increase precision when testing the interaction of THC and (*S*)-ketamine, we determined core body temperature every 30 min over a 90-min period using either 5 mg/kg THC or 10 mg/kg (*S*)-ketamine alone, or in combination (**Fig. 3b**). For THC, core body temperature decreased throughout the observation period (0.28 ± 0.20 °C [vehicle] *vs*. -1.35 ± 0.13 °C [THC, T = 30 min], *p* = 0.03; 0.18 ± 0.22 °C [vehicle] *vs*. -2.58 ± 0.39 °C [THC, T = 60 min], *p* = 0.01 and 0.8 ± 0.4 °C [vehicle] *vs*. -2.95 ± 0.62 °C [THC, T = 90 min]; *n* = 4-5), consistent with its long half-life and behavioral effects^26,45^.

After 30 min of 10 mg/kg (*S*)-ketamine exposure (T = 60 min), the core body temperature was reduced (-2.92 ± 0.46 °C; *p* = 0.01 *vs.* vehicle; *n* = 5) but improved after another 30 min (T = 90 min; -1.48 ± 1.19 °C; *p* = 0.2 *vs.* T = 60 min), in agreement with its short half-life^44^. When THC preceded (*S*)-ketamine, the core body temperature was significantly below of either drug alone, and failed to recover to pre-(*S*)-ketamine levels at either 60 or 90 min ([THC + (*S*)-ketamine, T = 60 min]: -4.80 ± 0.42°C and [THC + (*S*)-ketamine, T = 90 min]: -5.43 ± 0.48°C; *n* = 8), suggesting that (*S*)-ketamine augmented THC effects within this specific time window (**Fig. 3b**). Cumulatively, these data show that (*S*)-ketamine worsens pre-existing THC effects in a sequential setting.

### Serum availability of (*S*)-ketamine upon THC pretreatment

Drug interactions frequently intersect at the level of metabolic processes due to liver enzyme induction or inhibition^46^. As the metabolites of THC inhibit members of the cytochrome P450 family (CYP2B6, CYP2C9 and CYP2D6 in particular)^47^ and ketamine is degraded by CYP2B6 and CYP2C9^48^, we hypothesized that THC might alter the effects of (*S*)-ketamine by modulating its half-life in the circulation of mice. Nevertheless, the analysis of serum ketamine content by ELISA revealed no change in (*S*)- ketamine bioavailability in the presence of THC at 90 min, as compared to (*S*)-ketamine alone (**Fig. S6**). These data suggest that THC-(*S*)-ketamine interactions are unlikely to stem from the altered metabolism of (*S*)-ketamine.

### Stress hormones

THC increases both corticosterone and epinephrine levels by, e.g., inducing CRH release through type 1 cannabinoid receptors (CB_1_Rs) in the paraventricular nucleus (PVN) of the hypothalamus^49^, the uppermost module of the hypothalamus-pituitary-adrenal axis (HPA axis)^50^. In contrast, ketamine reduces circulating stress hormone levels, likely acting at (a) level(s) of the HPA axis where NMDAR subunits are expressed^31,32^. To test if THC and (*S*)-ketamine interact at the level of stress hormones, we collected truncal blood at 60 min and 120 min after THC exposure (that is, 30 min and 90 min after (*S*)-ketamine treatment), and first analyzed plasma adrenocorticotropic hormone (ACTH) content to mark pituitary activation^51^. After 60 min of THC exposure, ACTH levels tended to increase: ∼118% for THC, ∼163% for (*S*)-ketamine, and ∼146% for THC + (*S*)-ketamine, as compared to vehicle controls (normalized to 100%; **Fig. 3c**). Nevertheless, these data might be biased because the ACTH surge is fast, as is its feedback inhibition^52^, and might tail off differentially in drug-treated animals. Therefore, we determined corticosterone and epinephrine concentrations next, at both 60 min and 120 min post-THC exposure.

After 60 min of THC (5 mg/kg) exposure, we found increased corticosterone levels in plasma (∼380%; *p* < 0.001). (*S*)-ketamine (10 mg/kg) treatment alone also led to elevated circulating corticosterone levels (∼297%; *p* = 0.001 relative to controls; **Fig. 3d**). Combined treatment did not further boost corticosterone levels, likely due to a ‘ceiling effect’ at the drug concentrations used (∼347%; *p* < 0.001; **Fig. 3d**). After 120 min of THC exposure, corticosterone levels remained elevated (∼146%; *p* < 0.001; **Fig. 3d_1_**). In contrast, we noticed a significant reduction in corticosterone levels in mice treated with (*S*)-ketamine alone (∼65%; *p* = 0.02; **Fig. 3d_1_**). Nevertheless, corticosterone levels in the presence of both THC and (*S*)-ketamine remained significantly increased at this time point (∼142%; *p* = 0.004 *vs.* vehicle).

Lastly, we determined if epinephrine, released from the adrenal medulla by activation of the sympathetic nervous system upon stress, demonstrated a similar pattern. Indeed, epinephrine levels were significantly increase at 60 min in all drug-treated groups compared to vehicle (∼196% for THC; ∼175% for (*S*)-ketamine; ∼206% for THC + (*S*)-ketamine, *p* < 0.001 for all; **Fig. 3e**). Alike for corticosterone, (*S*)-ketamine effectively reduced epinephrine concentrations at 120 min (∼61%; *p* = 0.078; **Fig. 3e_1_**). However, combined treatment overrode the beneficial effect of (*S*)- ketamine alone (∼136%; *p* = 0.08 *vs.* vehicle and *p* < 0.001 *vs*. (*S*)-ketamine; **Fig. 3e_1_**). In sum, these data suggest that THC can offset the anxiolytic effects of (*S*)-ketamine, which can contribute to adverse drug interactions at the behavioral level.

### *Cnr1* and *Grin* mRNA expression in the HPA-axis

CB_1_Rs are expressed in the PVN, and also likely on corticotrope cells of the pituitary^53^ alike the adrenal medulla and cortex^54^, with their net effect (inhibition *vs*. stimulation) being dose- and stimulus severity-dependent. Similarly, NMDAR subunits show cell-type and tissue-specific distribution patterns. The co-expression of *Cnr1* and *Grin* mRNAs on cells in the HPA axis could thus be taken as a null hypothesis for drug interactions to occur. However, focused results specifying mRNA coexistence for CB_1_R and NMDARs in the HPA axis are as yet lacking. To this end, we reprocessed open-label RNA-seq datasets from the adult mouse hypothalamus^55^, pituitary and adrenal gland^56^. Both *Cnr1* and *Grin1*, *Grin2a*, *Grin2b* and *Grin3a* were expressed in *Crh*-containing neurons of the hypothalamus, including *Crh*^+^ PVN neurons (**Fig. 4a,a_1_**; *see also*^57^). The analysis of pituitary corticotropes, marked by *Pomc*, the precursor for ACTH^58^, revealed the unexpected lack of *Cnr1* (**Fig. 4b,b_1_**), with moderate expression of *Grin1*, *Grin2a*, and *Grin3a* mRNAs. Conversely, *Cyp11b1*-expressing cells of the adrenal cortex (noting that *Cyp11b1* encodes steroid-11β-hydroxylase, a rate-limiting enzyme for corticosterone biosynthesis^59^) primarily expressed *Cnr1* transcripts but not NMDAR subunits (**Fig. 4c,c_1_**). These data suggest that drug interactions could dominate at the level of CRH^+^ neurons^49^ in the HPA axis.

**Figure 4.**
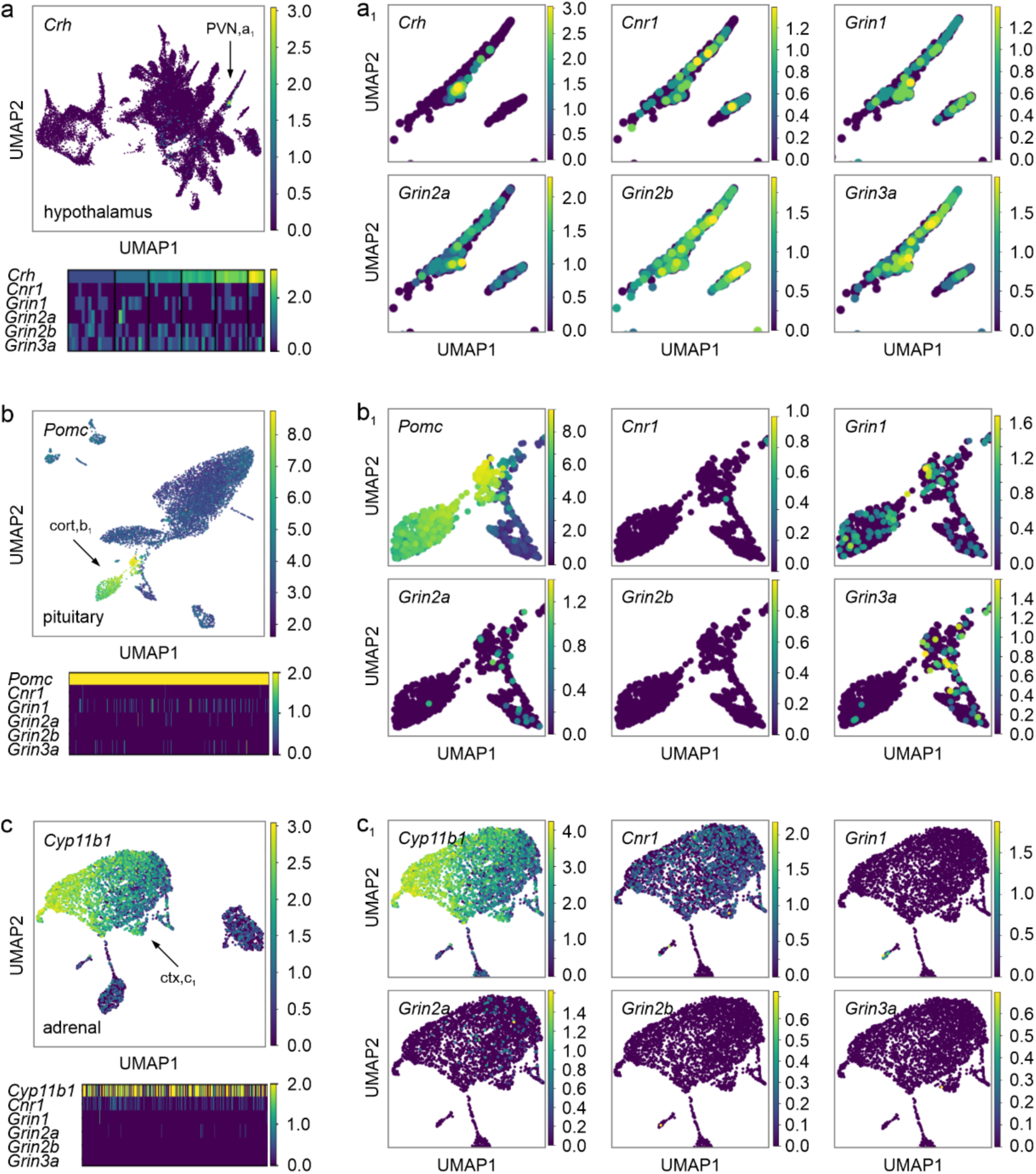
Single-cell RNA-seq-based co-expression mapping in the HPA axis. Publicly available datasets were reprocessed from Refs.^56,88^ (accession numbers: GSE132730 and GSE161751).(**a**) Uniform Manifold Approximation and Projection (UMAP; *top*) and a heat map (*bottom*) showing gene expression levels (from blue [minimum] to yellow [maximum]) for *Crh*, *Cnr1* and *Grin* genes (NMDAR subunits) in the hypothalamus. Arrow points to the UMAP position of paraventricular (PVN) *Crh*-containing neurons. (**a_1_**) Co-expression of all genes of interest in subpopulations of *Crh*-containing neurons (as per heat map), suggesting that both THC and (*S*)-ketamine could affect these neurons. (**b**) *Cnr1* and *Grin* mRNAs in pituitary corticotropes (cort) identified by *Pomc*, the ubiquitously expressed precursor to ACTH, in UMAP plots (*top*) and their target gene profiles (*bottom*). (**b_1_**) *Cnr1* mRNA was rarely, if at all, detected in *Pomc*- containing cells (UMAP insets for subpopulation specificity). In contrast, *Grin1*, *Grin2*, and *Grin3a* mRNAs were observed. (**c**) Cells of the adrenal cortex (ctx) harbored *Cyp11b1*, the rate-limiting enzyme for corticosterone synthesis (*top*). Target gene expression in *Cyp11b1*-expressing cells is at the bottom. (**c_1_**) *Cnr1* but not *Grin* family transcripts were abundant in *Cyp11b1*-expressing cells.

## Discussion

Polypharmacy, the use of two or more pharmaceuticals to treat multiple comorbid symptoms, is a current treatment standard in psychiatry^60^. For instance, patients suffering from clinical depression usually present multiple symptoms, including low mood, lack of motivation, anxiety and/or sleep disturbances^61^. Therefore, clinicians frequently co-prescribe benzodiazepines for anxiety, lamotrigine/lithium for mood alterations, as well as antipsychotics for sleep, in addition to occasional multiple antidepressant types for maximal health benefits^60^. Thus, it is becoming mandatory to map potential drug interactions to not only ensure appropriate pharmaceutical efficacy but also avoid life-long drug dependency. This is particularly valid for (*S*)-ketamine therapy, as it is commonly prescribed in addition to treatment regimens containing multiple medications for TRD and PTSD^62,63^. In addition, the increasing social acceptance of psychiatric treatment and ease of obtaining pharmaceuticals through online healthcare platforms^17^ spurred an upsurge in both professional and personal self-medication strategies. This not only led to beneficial personalized treatment protocols, but also to complications with misdiagnosis, inappropriate drug dosages, and treatment durations, as well as unexpected drug interactions^64^. The latter is highly relevant for recreational substance use, as drug liberalization is increasing worldwide^65,66^, production and availability have increased significantly throughout the last decade^67^, and their consumption remains growing over time^16,19^. While recent literature has focused on the interaction of licensed drugs with (*S*)-ketamine, data on interactions with recreational drugs, including cannabis, are lacking. Currently, the only indication of an interaction effect with cannabis is through one of its non-psychoactive phytocannabinoids, cannabidiol (CBD). Co-administration of CBD with (*S*)- ketamine led to an α-amino-3-hydroxy-5-methyl-4-isoxazolepropionic acid (AMPA)-dependent reduction in the hypermobility phase of acute ketamine exposure, without reducing its anti-depressant effects^68^. However, while CBD might be a beneficial adjunct therapy, the interaction with the major psychoactive phytocannabinoid, THC, remains unstudied. Thus, major findings of the present report are that (*S*)-ketamine-THC interactions are sequence specific, with (*S*)-ketamine added on top of THC leading to worsened behavioral, physiological, and endocrine outcomes.

Due to the complexity of dissecting the anti-depressant effects of pharmaceutical drugs in rodents with additional behavioral experiments^78^, especially in healthy animals, we opted to verify our findings by analyzing biomarkers associated with mental health disorders. For instance, activity of the HPA-axis^79^, and consequently the levels of circulating corticosterone (cortisol in humans), are directly associated with the severity of mental disorders, in particular depression and suicidal ideation^80^. We therefore analyzed both ACTH (pituitary) and corticosterone (adrenal cortex) in blood plasma. We further tested epinephrine, released from the adrenal medulla upon activation of the sympathetic nervous system, as it is implicated in patients with depressive symptoms^81^. While corticosterone and epinephrine levels proved relatively stable within treatment groups, ACTH measurements demonstrated wider inter-subject variation, even though its half-life is longer than of either corticosterone or epinephrine^82^. Despite the more variable results, both THC and (*S*)-ketamine increased ACTH after 30 min post (*S*)-ketamine exposure, suggesting pituitary activation. This was confirmed by the detection of increased circulating levels of corticosterone, substantiating the activation of the entire HPA axis. While the long term effects of (*S*)-ketamine led to a reduction in epinephrine and corticosterone, the presence of THC prevented a decrease of both hormones, suggesting that THC can overrule the therapeutic stress-regulatory effects of (*S*)-ketamine. We attribute this to the expression of CB_1_Rs across the anatomical constituents of the HPA axis overarching multiple organs. Specifically, endogenous cannabinoids (endocannabinoids) signal through CB_1_Rs in the paraventricular nucleus of the hypothalamus to tightly control CRH availability (**Supporting Fig. 7**). However, THC, as a relatively weak partial agonist at CB_1_Rs, is thought to disrupt endocannabinoid signaling by acting as a functional antagonist resulting in disinhibition of CRH-containing neurons, allowing for the downstream activation of the HPA axis, and ultimately increasing ACTH and then corticosterone availability^49^. As we find high levels of CB_1_Rs and NMDAR subunits expressed in the PVN by single-cell RNA-seq*^57^*, in contrast to the lack of CB_1_Rs in the pituitary and minor amounts of NMDARs in the adrenal cortex^56^, we hypothesize that the majority of interactions between THC and (*S*)-ketamine could occur in the hypothalamic locus of the HPA axis. However, a pituitary contribution can perhaps be expected as *in vitro* co-stimulation of cultured corticotrope cells with CRH and the cannabinoid receptor agonist WIN 55,212-2, but not with either substance alone, resulted in CB_1_R-dependent ACTH secretion^51^. These findings are particularly important, as the effects of THC can be maintained for considerable amounts of time due to its long half-life (20-30 h), with some of its active metabolites reaching half-lives of days to weeks^26^. Thus, exposure to THC could lead to longterm CB_1_R modulation impinging on the HPA axis to keep circulating corticosterone levels elevated^74^. Indeed, both acute and chronic cannabis users display higher levels of salivary cortisol and present HPA axis dysregulation^75,76^, which could offset the therapeutic reduction in stress hormones induced by (*S*)-ketamine treatment. In sum, and based on our rodent findings, we not only warn against simultaneous consumption of THC/cannabis and ketamine in recreational settings due to substantial adverse interactions but also propose to avoid the exposure to THC-containing products days before clinical (*S*)-ketamine sessions, to prevent blunting of its therapeutic benefits on stress hormone regulation.

## Acknowledgments

The authors thank Alexandra Wolf for her expert assistance. This work was supported by funding from the Austrian Science Fund FWF (P 34121-B, E.K.; 10.55776/COE16, T.H. and PAT7123823 (EFP9) to R.A.R.), Medical Neuroscience Cluster of the Medical University of Vienna (2023, E.K.), Vetenskapsrådet (2023-03058, T.H.), Novo Nordisk Foundation (NNF23OC0084476, T.H.), and the European Research Council (FOODFORLIFE, ERC-2020-AdG-101021016, T.H.).

## Data availability

ScRNA-seq datasets were curated from GEO and are available with accession codes: GSE132730 and GSE161751.

## Materials and methods

### Animals and ethical considerations

Male C57BL/6JRj mice 8-12 weeks of age (*n* = 174; Janvier) were group-housed conventionally (12h/12h light/dark cycle, 22-24 °C ambient temperature, and 55% humidity), and allowed ≥7-days of recovery upon arrival to reduce the impact of transport- and environmental change-related stress. Food and water were provided *ad libitum*. The sizes of animal groups were specified for each experiment separately. Animals were randomized, when necessary, yet without blinding. Sex differences were not addressed. Experiments were approved by the Austrian Ministry for Science and Research (GZ: 2025-0.844.019), and conformed to the European Communities Council Directive on the care and use of laboratory animals (2010/63/EU). Effort was directed towards minimizing the number of animals used and their suffering during the experiments. Group sizes conformed to those in the literature^27^. Animals were excluded only if they did not react to the drug(s) injected due to complications with drug delivery (with a maximum of *n* = 1 per experiment after THC exposure).

### Drug preparation and experimental use

Synthetic THC (Dronabinol; Gatt-Koller) stock solutions were prepared at a concentration of 62.5 mg/ml in dimethyl sulfoxide (DMSO; Sigma-Aldrich), aliquoted, and stored at −20 °C until use. Working solutions were freshly prepared daily by mixing 2% of the THC stock with 10% Cremophor (Kolliphor) EL (Sigma-Aldrich) and 88% sterile physiological saline (Braun) warmed to 37°C. Eskelan ((*S*)-ketamine, GL Pharma, 25 mg/ml) was diluted in sterile physiological saline (Braun) daily. For both compounds, vehicle solutions were prepared using the same formulations excluding the active ingredient. Final preparations were vortexed before use. For all injections, Omnifix-F Luer Lock Solo syringes (0.01– 1 mL; Braun) were fitted with Sterican needles (0.30 × 12 mm; Braun) to inject ± 100 µl of compound intraperitoneally, with the final volume adjusted to the weight of the experimental subjects. Injections sites were alternated between the left *vs*. right abdomen for successive injections.

### Determination of core body temperature

Core body temperature was measured in 30-min intervals before and after drug infusion by using a rectal thermoprobe (Physitemp BAT-10) specifically designed for small laboratory animals. The Vaseline-coated probe was inserted approximately 1 cm into the clean recta of the experimental subjects to ensure consistent and reliable readings across all animals.

### Behavioral tests

On the day of the experiments, the bodyweights of the mice were first measured (baseline) after which they were returned to their cages in the behavioral room for ≥60 min. Mice were randomly assigned to the experimental groups and members of each group were tested between 9:00 and 15:00 on experimental days. Ambulation, exploratory and maintenance behaviors were determined in an open field consisting of a rectangular arena (42 x 42 x 21 cm with white background) with an overhead-centered artificial light source (light intensity: 210-220 Lux for all of the experiments; Voltcraft LX-10). The arena was cleaned with 70% ethanol and thoroughly dried after each experiment. At the start, mice were placed in the center of the arena with their activity recorded for 30 min afterwards. Videos were analyzed offline by Ethovision (17.0, Noldus) with parameters including: distance moved (cm), grooming (% of total time), unsupported rearing (% of total time), supported rearing (% of total time), and rotations around the body axis (number of rotations per meter travelled).

Motor coordination and balance were assessed using a rotarod (Ugo Basile, Accelerating Rota-Rod). Mice were familiarized with the test apparatus by repeatedly placing them on the stationary rotarod, thus also reducing novelty-induced stress. During the actual test, mice were placed on the rotating cylinder. The acceleration mode was used for all trials. Each animal completed three consecutive trials. A trial was considered complete when the mouse either fell off the rod or completed one passive rotation (i.e., clung to the rod without active locomotion). The latency to fall was recorded for each trial (s). The mean of the three trials was used as a measure of performance.

For the catalepsy test, mice were placed with their front paws on a horizontal bar measuring 3.5 cm above the cage floor^27^. For each trial, the time was measured until the mouse would leave the horizontal bar, for up to three trials. However, due to the strong startle response there were complications to reliably place the mice on the bar with THC concentrations ≥ 5mg/kg (*see Supporting movies S1,S2*).

### ELISA-based determination of serum ketamine and hormone levels

After the behavioral tests, mice were anesthetized by isoflurane (CP Pharma; 5% in 1L/min flow of tubed air) and decapitated. Truncal blood was collected, allowed to clot for 30 min at 22-24 °C, and then centrifuged at 1,200 *g* at 4 °C for 10 min. Serum was aspirated and stored at -80 °C until further use. Enzyme-linked immunosorbent assay (ELISA) kits for ketamine (Neogen, 131719), epinephrine (Abcam, AB287788), corticosterone (Enzo Biosciences, ADI-900-097) and ACTH (Abcam, AB263880) were used to determine their respective concentrations in diluted sera (1:20 for cortisol, 1:10 for ketamine, 1:10 for epinephrine, and 1:3 for ACTH). Data were normalized to the mean of vehicle-treated controls.

### Processing of single-cell RNA-seq

Open-access single-cell RNA-seq datasets for the adult mouse hypothalamus^88^, pituitary and adrenal cortex^56^ with accession numbers GSE132730 and GCE161751 were subset for *Crh* among hypothalamic PVN neurons^57^; *Pomc* for pituitary corticotropes^58^, and *Cyp11b1* for cortisol-producing cells of the adrenal cortex^59^. Subsequently, pairwise co-expression of cell typifying markers and *Cnr1* and *Grin*-family mRNAs (the latter encoding NMDAR subunits) were processed. For the pituitary and adrenal data, cells with low gene counts (<1,000) or high mitochondrial transcript content (>20%) were excluded. Data were normalized to the median total counts per cell and log transformed. Analyses were performed in Python using *Scanpy* (version 1.9.5). Highly variable genes were selected using the *Scanpy* default method. Principal component analysis (PCA) was performed, and the first 30 principal components were used to construct a neighborhood graph. Uniform Manifold Approximation and Projection (UMAP) was applied for two-dimensional visualization, with clusters identified using the Leiden algorithm with a resolution of 0.5. For the hypothalamic dataset, *geneset* visualization was performed with the Seurat object (GSE132730_subset_cca_to_velo.rds.gz) and processed as previously described^57,88^. Data were plotted as co-expression matrices (using SCT-transformed counts) and UMAP representations using routine analysis pipelines^57,88^.

### Statistics

Statistical evaluation was performed in GraphPad Prism (v.10.5.0) with one-way ANOVA (for multifactorial testing, followed by an uncorrected LSD or Tukey test), Kruskal-Wallis tests (for non-normal data distribution), or two-trailed Student’s *t*-test (normal distribution) depending on the null hypothesis queried. Data were expressed as means ± s.e.m. A *p* value of <0.05 was considered statistically significant. All data and statistical outcomes for each experiment presented were uploaded as a separate Source Data file (*.Xlsx*).

## References

1. Kasper, S. et al. Practical recommendations for the management of treatment-resistant depression with esketamine nasal spray therapy: Basic science, evidence-based knowledge and expert guidance. The World Journal of Biological Psychiatry 22, 468–482 (2021).

2. Hillhouse, T. M. & Porter, J. H. A brief history of the development of antidepressant drugs: From monoamines to glutamate. Exp Clin Psychopharmacol 23, 1–21 (2015).

3. Kumari, S. et al. Exploring Esketamine’s Therapeutic Outcomes as an FDA-Designated Breakthrough for Treatment-Resistant Depression and Major Depressive Disorder With Suicidal Intent: A Narrative Review. Cureus 16, e53987 (2024).

4. McIntyre, R. S. et al. Synthesizing the Evidence for Ketamine and Esketamine in Treatment-Resistant Depression: An International Expert Opinion on the Available Evidence and Implementation. Am J Psychiatry 178, 383–399 (2021).

5. Zarate, C. A. et al. A randomized trial of an N-methyl-D-aspartate antagonist in treatment-resistant major depression. Arch Gen Psychiatry 63, 856–864 (2006).

6. Berman, R. M. et al. Antidepressant effects of ketamine in depressed patients. Biological Psychiatry 47, 351–354 (2000).

7. Medeiros, G. C., Demo, I., Goes, F. S., Zarate, C. A. & Gould, T. D. Personalized use of ketamine and esketamine for treatment-resistant depression. Transl Psychiatry 14, 481 (2024).

8. Kritzer, M. D. et al. Ketamine for treatment of mood disorders and suicidality: A narrative review of recent progress. Ann Clin Psychiatry 34, 33–43 (2022).

9. SPRAVATO® (esketamine). SPRAVATO® (esketamine) | Official Patient Website https://www.spravato.com/ (2025).

10. Fan, N. et al. Profiling the psychotic, depressive and anxiety symptoms in chronic ketamine users. Psychiatry Res 237, 311–315 (2016).

11. Nikayin, S., Murphy, E., Krystal, J. H. & Wilkinson, S. T. Long-term safety of ketamine and esketamine in treatment of depression. Expert Opin Drug Saf 21, 777–787 (2022).

12. Strous, J. F. M. et al. Brain Changes Associated With Long-Term Ketamine Abuse, A Systematic Review. Front Neuroanat 16, 795231 (2022).

13. Bavato, F. & Quednow, B. B. Ketamine addiction in Europe: Any risks on the horizons? European Neuropsychopharmacology 94, 39–40 (2025).

14. Coplan, J. D., Aaronson, C. J., Panthangi, V. & Kim, Y. Treating comorbid anxiety and depression: Psychosocial and pharmacological approaches. World J Psychiatry 5, 366–378 (2015).

15. Veraart, J. K. E. et al. Pharmacodynamic Interactions Between Ketamine and Psychiatric Medications Used in the Treatment of Depression: A Systematic Review. Int J Neuropsychopharmacol 24, 808–831 (2021).

16. EUDA. EU Drug Market: New psychoactive substances — Distribution and supply in Europe: Ketamine. https://www.euda.europa.eu/publications/eu-drug-markets/new-psychoactive-substances/distribution-and-supply/ketamine_en.

17. Hull, T. D. et al. At-home, sublingual ketamine telehealth is a safe and effective treatment for moderate to severe anxiety and depression: Findings from a large, prospective, open-label effectiveness trial. J Affect Disord 314, 59–67 (2022).

18. Lankenau, S. E. & Clatts, M. C. Patterns of polydrug use among ketamine injectors in New York City. Subst Use Misuse 40, 1381–1397 (2005).

19. EUDA. Cannabis – the current situation in Europe (European Drug Report 2025). https://www.euda.europa.eu/publications/european-drug-report/2025/cannabis_en.

20. Freeman, T. P. et al. Changes in delta-9-tetrahydrocannabinol (THC) and cannabidiol (CBD) concentrations in cannabis over time: systematic review and meta-analysis. Addiction 116, 1000–1010 (2021).

21. Beletsky, A., Liu, C., Lochte, B., Samuel, N. & Grant, I. Cannabis and Anxiety: A Critical Review. Med Cannabis Cannabinoids 7, 19–30 (2024).

22. Li, X. et al. The Effectiveness of Cannabis Flower for Immediate Relief from Symptoms of Depression. Yale J Biol Med 93, 251–264 (2020).

23. Audet, C. et al. Self-Medication Paths: A Descriptive Study Unveiling the Interplay Between Medical and Nonmedical Cannabis in Chronic Pain Management. Clin J Pain 40, 635–645 (2024).

24. Fletcher, P. C. & Honey, G. D. Schizophrenia, ketamine and cannabis: evidence of overlapping memory deficits. Trends in Cognitive Sciences 10, 167–174 (2006).

25. Matsuda, L. A., Lolait, S. J., Brownstein, M. J., Young, A. C. & Bonner, T. I. Structure of a cannabinoid receptor and functional expression of the cloned cDNA. Nature 346, 561–564 (1990).

26. Wall, M. E. & Perez-Reyes, M. The metabolism of delta 9-tetrahydrocannabinol and related cannabinoids in man. J Clin Pharmacol 21, 178S–189S (1981).

27. Metna-Laurent, M., Mondésir, M., Grel, A., Vallée, M. & Piazza, P.-V. Cannabinoid-Induced Tetrad in Mice. Current Protocols in Neuroscience 80, 9.59.1–9.59.10 (2017).

28. Irifune, M., Shimizu, T. & Nomoto, M. Ketamine-induced hyperlocomotion associated with alteration of presynaptic components of dopamine neurons in the nucleus accumbens of mice. Pharmacology Biochemistry and Behavior 40, 399–407 (1991).

29. Zaytseva, A. et al. Ketamine’s rapid antidepressant effects are mediated by Ca2+-permeable AMPA receptors. eLife 12, e86022 (2023).

30. Simmler, L. D. et al. Dual action of ketamine confines addiction liability. Nature 608, 368–373 (2022).

31. Dutton, M., Can, A. T., Lagopoulos, J. & Hermens, D. F. Stress, mental disorder and ketamine as a novel, rapid acting treatment. European Neuropsychopharmacology 65, 15–29 (2022).

32. Ostroff, R. & Kothari, J. S. Reversal of Non-Suppression of Cortisol Levels in a Patient With Refractory Depression Receiving Ketamine. AJP 172, 95–96 (2015).

33. Lulek, C. F. et al. Sex differences in acute delta-9-tetrahydrocannabinol (Δ9-THC) response and tolerance as a function of mouse strain. Psychopharmacology (Berl) 240, 1987–2003 (2023).

34. Harda, Z., Misiołek, K., Klimczak, M., Chrószcz, M. & Rodriguez Parkitna, J. C57BL/6N mice show a sub-strain specific resistance to the psychotomimetic effects of ketamine. Front Behav Neurosci 16, 1057319 (2022).

35. Tannenbaum, C. & Day, D. Age and sex in drug development and testing for adults. Pharmacological Research 121, 83–93 (2017).

36. Nair, A. B. & Jacob, S. A simple practice guide for dose conversion between animals and human. J Basic Clin Pharm 7, 27–31 (2016).

37. Stival, C. et al. Prevalence and Correlates of Overweight and Obesity in 12 European Countries in 2017–2018. Obes Facts 15, 655–665 (2022).

38. Andrade, C. Ketamine for Depression, 4: In What Dose, at What Rate, by What Route, for How Long, and at What Frequency? J Clin Psychiatry 78, e852–e857 (2017).

39. Lin, M. T., Chen, C. F. & Pang, I. H. Effect of ketamine on thermoregulation in rats. Can J Physiol Pharmacol 56, 963–967 (1978).

40. Galvanho, J. P. et al. Profiling of behavioral effects evoked by ketamine and the role of 5HT2 and D2 receptors in ketamine-induced locomotor sensitization in mice. Prog Neuropsychopharmacol Biol Psychiatry 97, 109775 (2020).

41. Myslobodsky, M. S., Ackermann, R. F., Golovchinsky, V. & Engel, J. Ketamine-induced rotation: Interaction with GABA-transaminase inhibitors and picrotoxin. Pharmacology Biochemistry and Behavior 11, 483–486 (1979).

42. Sharma, P., Murthy, P. & Bharath, M. M. S. Chemistry, Metabolism, and Toxicology of Cannabis: Clinical Implications. Iran J Psychiatry 7, 149–156 (2012).

43. Blake, A. & Nahtigal, I. The evolving landscape of cannabis edibles. Current Opinion in Food Science 28, 25–31 (2019).

44. Ballard, E. D. & Zarate, C. A. The role of dissociation in ketamine’s antidepressant effects. Nat Commun 11, 6431 (2020).

45. Britch, S. C., Wiley, J. L., Yu, Z., Clowers, B. H. & Craft, R. M. Cannabidiol-Δ9-tetrahydrocannabinol interactions on acute pain and locomotor activity. Drug and Alcohol Dependence 175, 187–197 (2017).

46. Tanaka, E. Clinically important pharmacokinetic drug-drug interactions: role of cytochrome P450 enzymes. J Clin Pharm Ther 23, 403–416 (1998).

47. Nasrin, S., Watson, C. J. W., Perez-Paramo, Y. X. & Lazarus, P. Cannabinoid Metabolites as Inhibitors of Major Hepatic CYP450 Enzymes, with Implications for Cannabis-Drug Interactions. Drug Metab Dispos 49, 1070–1080 (2021).

48. Yanagihara, Y. et al. Involvement of CYP2B6 in n-demethylation of ketamine in human liver microsomes. Drug Metab Dispos 29, 887–890 (2001).

49. Hillard, C. J., Beatka, M. & Sarvaideo, J. Endocannabinoid Signaling and the Hypothalamic-Pituitary-Adrenal Axis. Compr Physiol 7, 1–15 (2016).

50. Russell, G. & Lightman, S. The human stress response. Nat Rev Endocrinol 15, 525–534 (2019).

51. Aguilera, G. Regulation of pituitary ACTH secretion during chronic stress. Front Neuroendocrinol 15, 321–350 (1994).

52. Osterlund, C. D. et al. Glucocorticoid Fast Feedback Inhibition of Stress-Induced ACTH Secretion in the Male Rat: Rate Independence and Stress-State Resistance. Endocrinology 157, 2785–2798 (2016).

53. Pagotto, U. et al. Normal human pituitary gland and pituitary adenomas express cannabinoid receptor type 1 and synthesize endogenous cannabinoids: first evidence for a direct role of cannabinoids on hormone modulation at the human pituitary level. J Clin Endocrinol Metab 86, 2687–2696 (2001).

54. Ziegler, C. G. et al. Expression and function of endocannabinoid receptors in the human adrenal cortex. Horm Metab Res 42, 88–92 (2010).

55. Romanov, R. A. et al. Molecular interrogation of hypothalamic organization reveals distinct dopamine neuronal subtypes. Nat. Neurosci. 20, 176–188 (2017).

56. Lopez, J. P. et al. Single-cell molecular profiling of all three components of the HPA axis reveals adrenal ABCB1 as a regulator of stress adaptation. Sci Adv 7, eabe4497 (2021).

57. Tretiakov, E. O. et al. Molecular Fingerprint of Endocannabinoid Signaling in the Developing Paraventricular Nucleus of the Hypothalamus as Revealed by Single-Cell RNA-Seq and In Situ Hybridization. Cells 14, 788 (2025).

58. Raffin-Sanson, M. L., de Keyzer, Y. & Bertagna, X. Proopiomelanocortin, a polypeptide precursor with multiple functions: from physiology to pathological conditions. Eur J Endocrinol 149, 79–90 (2003).

59. Miller, W. L. & Auchus, R. J. The Molecular Biology, Biochemistry, and Physiology of Human Steroidogenesis and Its Disorders. Endocr Rev 32, 81–151 (2011).

60. Wiersema, C. et al. Determinants and consequences of polypharmacy in patients with a depressive disorder in later life. Acta Psychiatr Scand 146, 85–97 (2022).

61. Thaipisuttikul, P., Ittasakul, P., Waleeprakhon, P., Wisajun, P. & Jullagate, S. Psychiatric comorbidities in patients with major depressive disorder. Neuropsychiatr Dis Treat 10, 2097–2103 (2014).

62. McIntyre, R. S. et al. Treatment-resistant depression: definition, prevalence, detection, management, and investigational interventions. World Psychiatry 22, 394–412 (2023).

63. Almeida, T. M. et al. Effectiveness of Ketamine for the Treatment of Post-Traumatic Stress Disorder – A Systematic Review and Meta-Analysis. Clin Neuropsychiatry 21, 22–31.

64. Hughes, C. M., McElnay, J. C. & Fleming, G. F. Benefits and risks of self medication. Drug Saf 24, 1027–1037 (2001).

65. Holland, A., Etches, S. & Gander, S. Drug decriminalization: The importance of policy change for the health and wellbeing of children and youth in Canada. Paediatr Child Health 29, 87–89 (2023).

66. Moury, C. & Escada, M. Understanding successful policy innovation: The case of Portuguese drug policy. Addiction 118, 967–978 (2023).

67. EUDA. Drug supply, production and precursors – the current situation in Europe (European Drug Report 2025) | www.euda.europa.eu. https://www.euda.europa.eu/publications/european-drug-report/2025/drug-supply-production-and-precursors_en.

68. Sartim, A. G. et al. Co-administration of cannabidiol and ketamine induces antidepressant-like effects devoid of hyperlocomotor side-effects. Neuropharmacology 195, 108679 (2021).

69. Bartova, L. et al. Rapid antidepressant effect of S-ketamine in schizophrenia. Eur Neuropsychopharmacol 28, 980–982 (2018).

70. Ghosal, S. et al. Ketamine rapidly reverses stress-induced impairments in GABAergic transmission in the prefrontal cortex in male rodents. Neurobiology of Disease 134, 104669 (2020).

71. Kokkinou, M., Ashok, A. H. & Howes, O. D. The effects of ketamine on dopaminergic function: meta-analysis and review of the implications for neuropsychiatric disorders. Mol Psychiatry 23, 59–69 (2018).

72. Han, X. et al. Cannabinoid CB1 Receptors Are Expressed in a Subset of Dopamine Neurons and Underlie Cannabinoid-Induced Aversion, Hypoactivity, and Anxiolytic Effects in Mice. J Neurosci 43, 373–385 (2023).

73. Bloomfield, M. A. P., Ashok, A. H., Volkow, N. D. & Howes, O. D. The effects of Δ9-tetrahydrocannabinol on the dopamine system. Nature 539, 369–377 (2016).

74. Rigucci, S. et al. Cannabis use in early psychosis is associated with reduced glutamate levels in the prefrontal cortex. Psychopharmacology (Berl) 235, 13–22 (2018).

75. Hwang, E.-K. & Lupica, C. R. Altered Corticolimbic Control of the Nucleus Accumbens by Longterm Δ9-Tetrahydrocannabinol Exposure. Biological Psychiatry 87, 619–631 (2020).

76. Hungund, B. L. et al. Upregulation of CB1 receptors and agonist-stimulated [35S]GTPgammaS binding in the prefrontal cortex of depressed suicide victims. Mol Psychiatry 9, 184–190 (2004).

77. Wang, W. et al. Deficiency in endocannabinoid signaling in the nucleus accumbens induced by chronic unpredictable stress. Neuropsychopharmacology 35, 2249–2261 (2010).

78. Planchez, B., Surget, A. & Belzung, C. Animal models of major depression: drawbacks and challenges. J Neural Transm (Vienna) 126, 1383–1408 (2019).

79. García-Gutiérrez, M. S. et al. Biomarkers in Psychiatry: Concept, Definition, Types and Relevance to the Clinical Reality. Front Psychiatry 11, 432 (2020).

80. Dziurkowska, E. & Wesolowski, M. Cortisol as a Biomarker of Mental Disorder Severity. J Clin Med 10, 5204 (2021).

81. Gold, S. M., Zakowski, S. G., Valdimarsdottir, H. B. & Bovbjerg, D. H. Higher Beck depression scores predict delayed epinephrine recovery after acute psychological stress independent of baseline levels of stress and mood. Biol Psychol 67, 261–273 (2004).

82. Veldhuis, J. D. et al. Corticotropin secretory dynamics in humans under low glucocorticoid feedback. J Clin Endocrinol Metab 86, 5554–5563 (2001).

83. Buckley, N. E., Hansson, S., Harta, G. & Mezey, E. Expression of the CB1 and CB2 receptor messenger RNAs during embryonic development in the rat. Neuroscience 82, 1131–1149 (1998).

84. Murphy, L. L., Muñoz, R. M., Adrian, B. A. & Villanúa, M. A. Function of cannabinoid receptors in the neuroendocrine regulation of hormone secretion. Neurobiol Dis 5, 432–446 (1998).

85. Cone, E. J., Johnson, R. E., Moore, J. D. & Roache, J. D. Acute effects of smoking marijuana on hormones, subjective effects and performance in male human subjects. Pharmacol Biochem Behav 24, 1749–1754 (1986).

86. Carol, E. E., Spencer, R. L. & Mittal, V. A. The relationship between cannabis use and cortisol levels in youth at ultra high-risk for psychosis. Psychoneuroendocrinology 83, 58–64 (2017).

87. al’Absi, M. & Allen, A. M. Impact of Acute and Chronic Cannabis Use on Stress Response Regulation: Challenging the Belief That Cannabis Is an Effective Method for Coping. Front. Psychol. 12, (2021).

88. Romanov, R. A. et al. Molecular design of hypothalamus development. Nature 1–7 (2020) doi:10.1038/s41586-020-2266-0.

