## Supporting Information File for "(*S*)-ketamine augments behavioral and physiological responses to Δ^9^-tetrahydrocannabinol"

**Contents:**

- Supporting figures
- Legends to supporting figures
- Legends to supporting movies
- Supporting references

### Supporting Figures

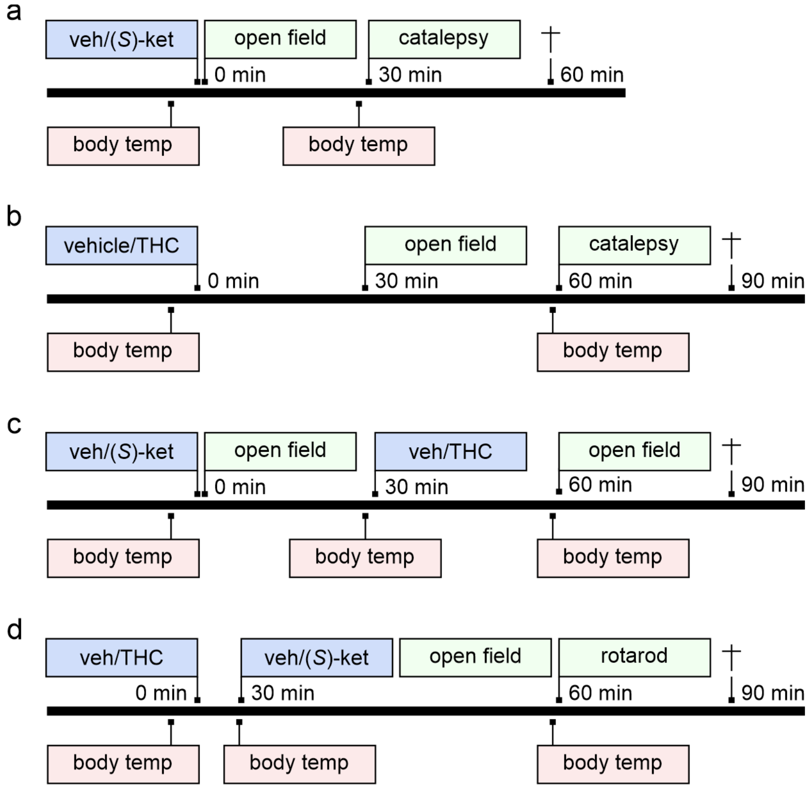


**Supporting Figure 1. Schema of the various experiments in the sequence they appear in the body text.** (**a,b**) Experimental layout to test dose-response relationships for either (*S*)-ketamine (**a**) or THC (**b**). (**c**) Experimental design to test drug interactions when (*S*)-ketamine treatment preceded that of THC. (**d**) Opposite sequence of drug application with THC prior (*S*)-ketamine.

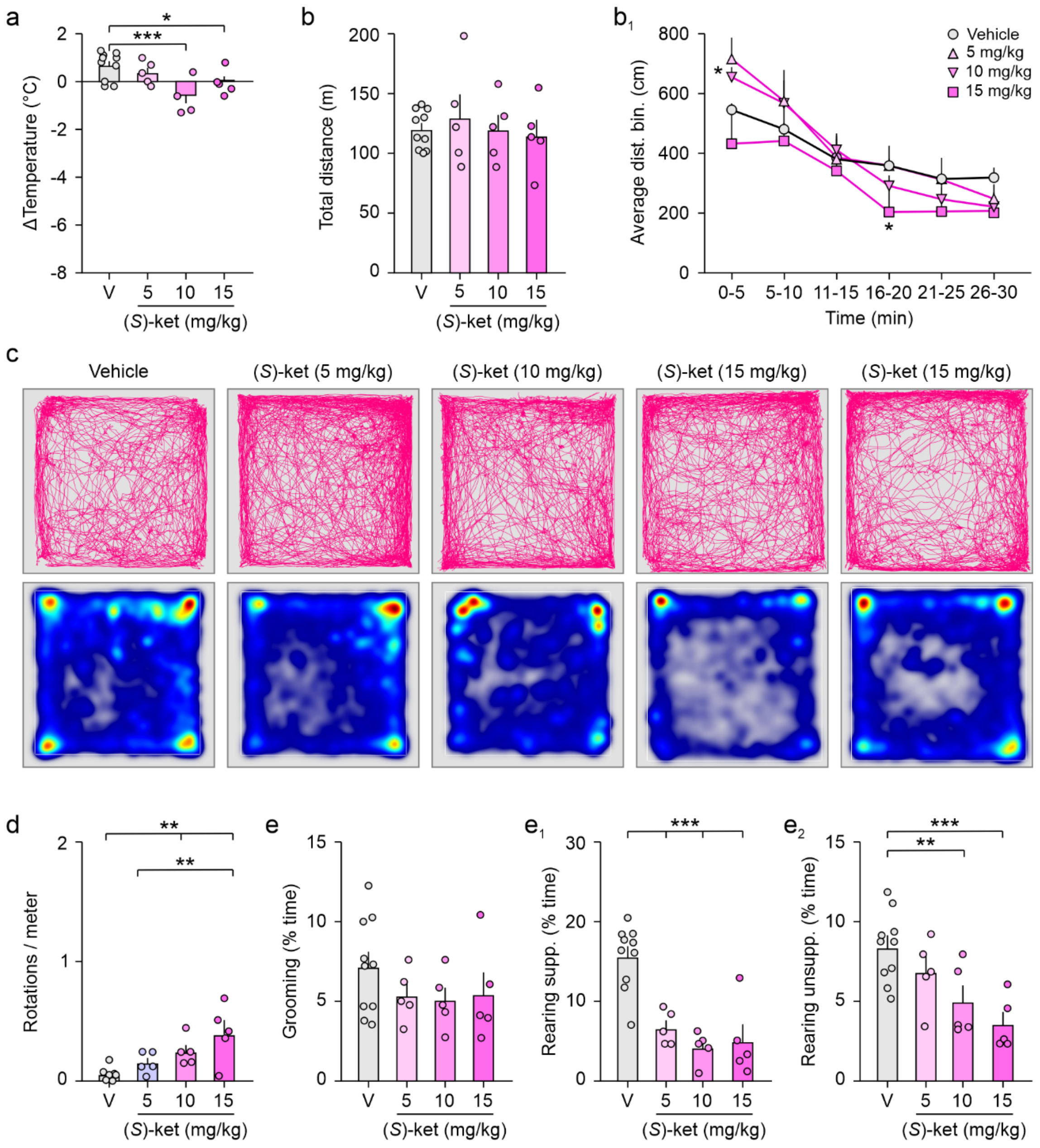


**Supporting Figure 2. Dose-response relationship for (*S*)-ketamine on core body temperature and exploratory behavior.** (**a**) Core body temperature 30 min after (*S*)-ketamine exposure. Cumulative (**b**) and binned (5 min, **b$_1$**) distances travelled in an open field in 30 min immediately after (*S*)-ketamine injection (*s.c.*). (**c**) Representative trajectories (*top*) and heat maps of time spent at specific locations in an open field (*bottom*; blue-to-red color scale refers to locations with none-to-maximum occupancy). (**d**) Number of axial rotations ('circling behavior'). (**e**-**e$_2$**) Grooming and rearing behavior in the open field. Data were aggregated over 30 min. Colored circles denote individual data points. Male mice were used throughout. Bar graphs show means ± s.e.m.; **p* < 0.05, ***p* < 0.01, ****p* < 0.001, ns, non-significant. *Abbreviations:* (*S*)-ket, (*S*)-ketamine at the concentrations shown in (mg/kg); V, vehicle.

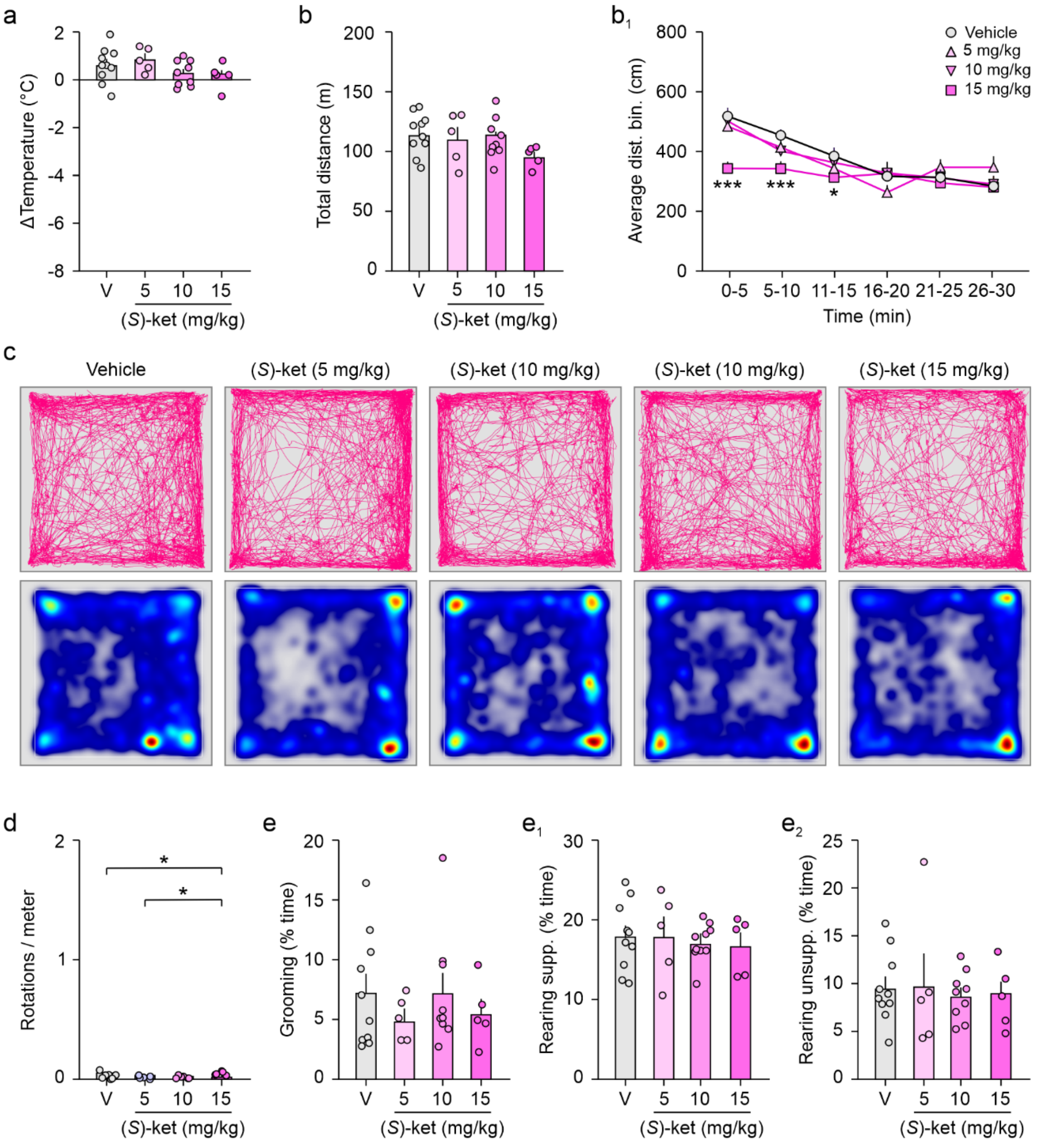


**Supporting Figure 3. Dose-response relationship for (*S*)-ketamine on core body temperature and exploratory behavior in a delayed test paradigm.** (**a**) Core body temperature 60 min after (*S*)-ketamine injection (*s.c.*, *n* = 5-10/group). Cumulative (**b**) and binned (5 min, $\mathbf{b_1}$) distances travelled in an open field for 30 min. Behavioral testing started 60 min after (*S*)-ketamine administration. (**c**) Representative trajectories (*top*) and heat maps of time spent at specific locations (*bottom*; blue-to-red color scale refers to locations with none-to-maximum occupancy). (**d**) Number of axial rotations ('circling behavior'). ($\mathbf{e}$-$\mathbf{e_2}$) Grooming and rearing in the open field. Data were aggregated over 30 min. Colored circles denote individual data points of male mice. Bar graphs show means ± s.e.m.; *$p < 0.05$, ***$p < 0.001$. *Abbreviations:* (*S*)-ket, (*S*)-ketamine at the concentrations shown in (mg/kg); V, vehicle.

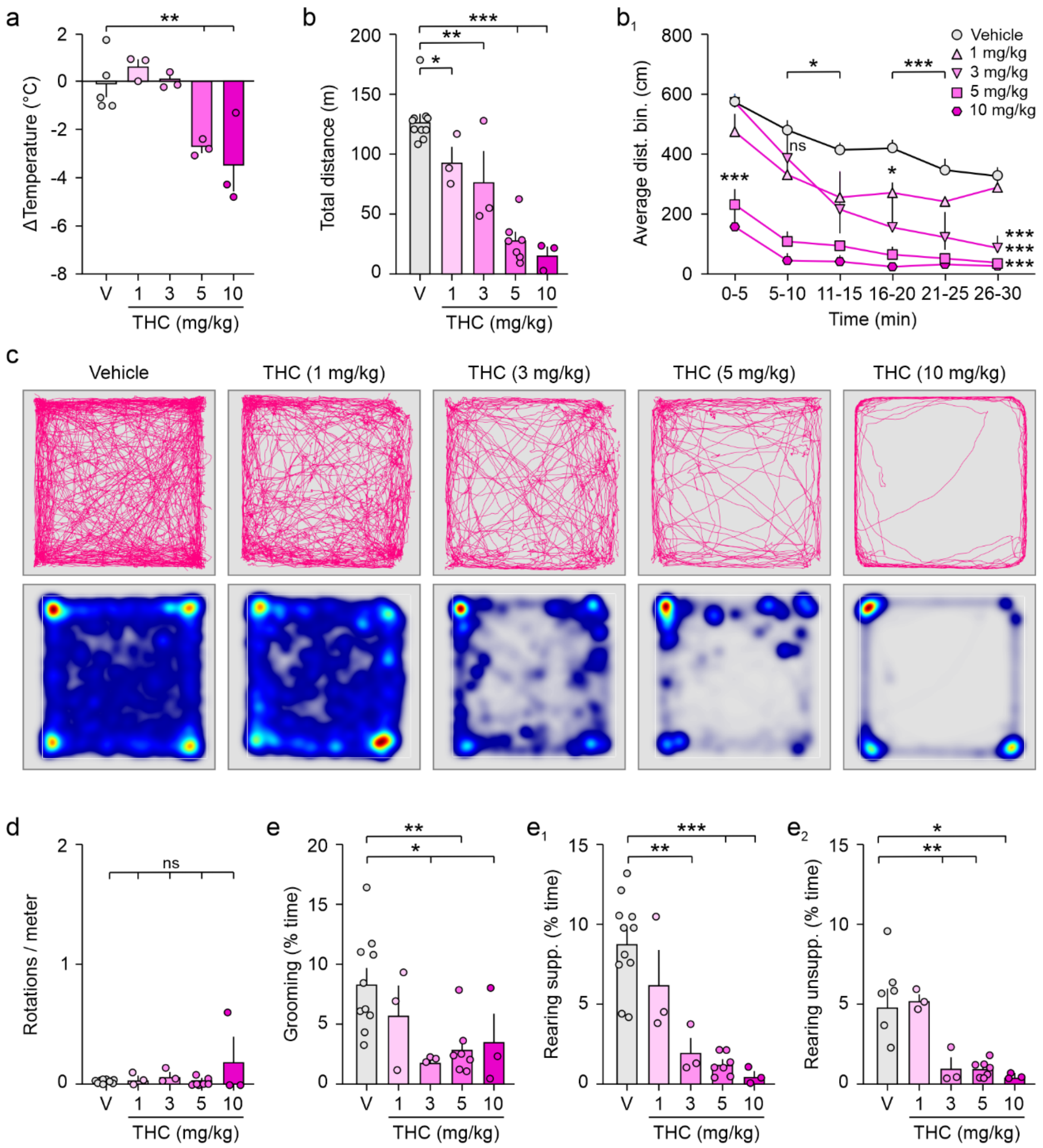


**Supporting Figure 4. Dose-response relationship for THC on core body temperature and exploratory behavior.** (**a**) Core body temperature 30 min after THC exposure (*s.c.*, $n = 3$-6/group). Cumulative (**b**) and binned (5 min, $\mathbf{b_1}$) distances travelled in an open field for 30 min. Behavioral testing started 30 min after THC injection. (**c**) Representative trajectories (*top*) and heat maps of time spent at specific locations (*bottom*; blue-to-red color scale refers to locations with none-to-maximum occupancy). (**d**) Number of axial rotations ('circling behavior'). ($\mathbf{e}$-$\mathbf{e_2}$) Grooming and rearing behavior in the open field accumulated in 30 min. Data were aggregated over 30 min. Colored circles denote individual data points of male mice. Bar graphs show means ± s.e.m. $*p < 0.05$, $**p < 0.01$, $***p < 0.001$, ns, non-significant. *Abbreviation:* V, vehicle.

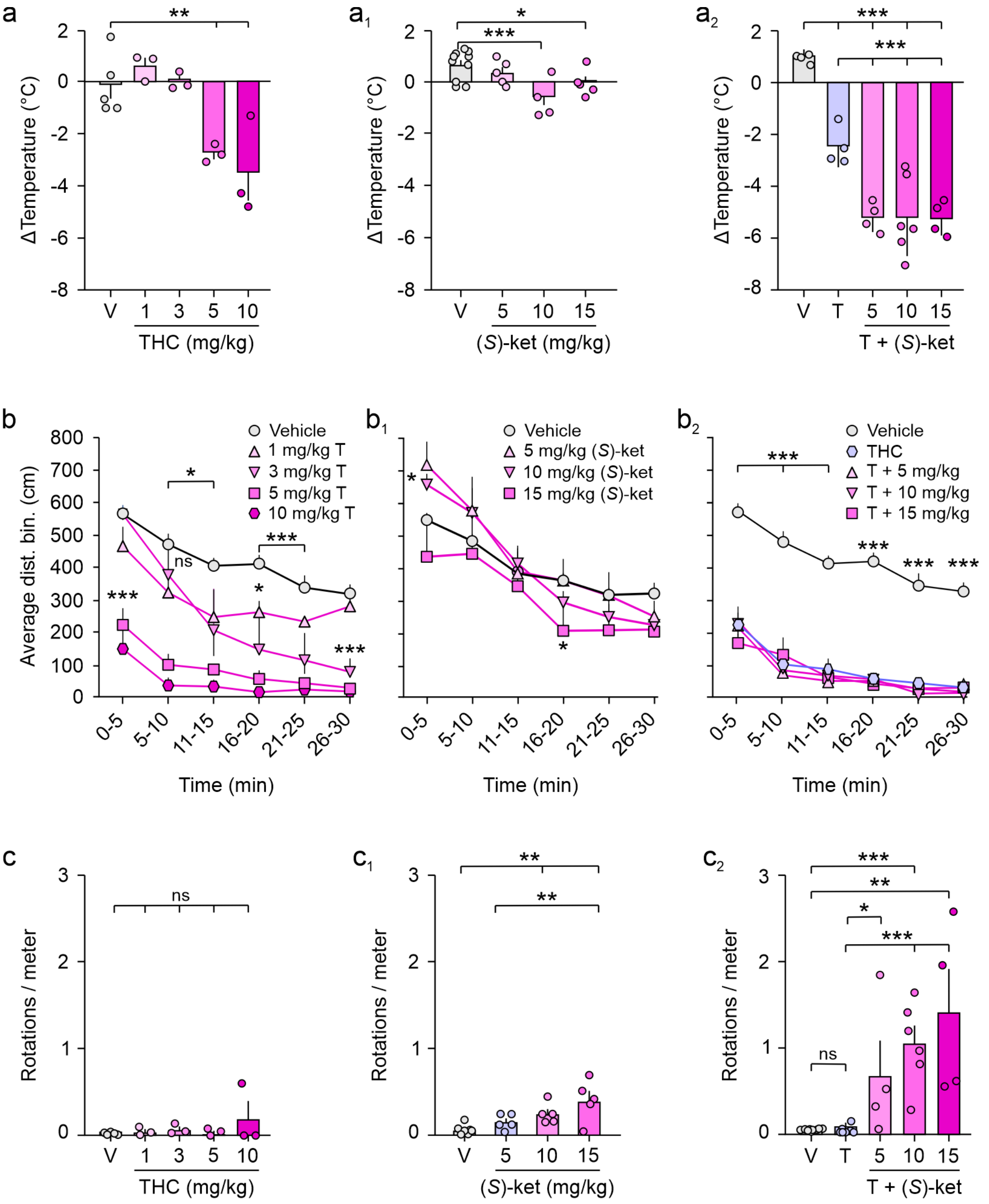


**Supporting Figure 5. Comparative experimental data for visual clarity.** (**a-c₂**) Data were scaled for temperature change (a-a₂), binned distance moved in the open field test ($n$ = 5-10 (**b**); $n$ = 3-10 (**b₁**); $n$ = 4-10 (**b₂**)), and the amount of circular rotations around the body axis (**c-c₂**). Colored circles denote individual data points of male mice. Bar graphs show means ± s.e.m.; *$p$ < 0.05, **$p$ < 0.01, ***$p$ < 0.001, ns, non-significant. *Abbreviations:* (*S*)-ket, (*S*)-ketamine; T, THC at the concentrations shown in (mg/kg); V, vehicle.

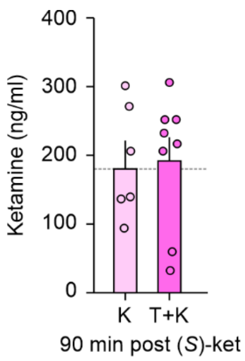


**Supporting Figure 6. Plasma serum concentrations of (*S*)-ketamine.** A forensic ELISA kit for ketamine (Neogen) was used to show that THC applied 30 min prior (*S*)-ketamine (T+K) did not alter serum ketamine concentrations at 90 min relative to (*S*)-ketamine use alone (K). Colored circles denote individual data points from male mice. Bar graphs show means ± s.e.m. Horizontal dashed line denotes the mean control value. *Abbreviations:* K, (*S*)-ketamine; T+K, THC + (*S*)-ketamine.

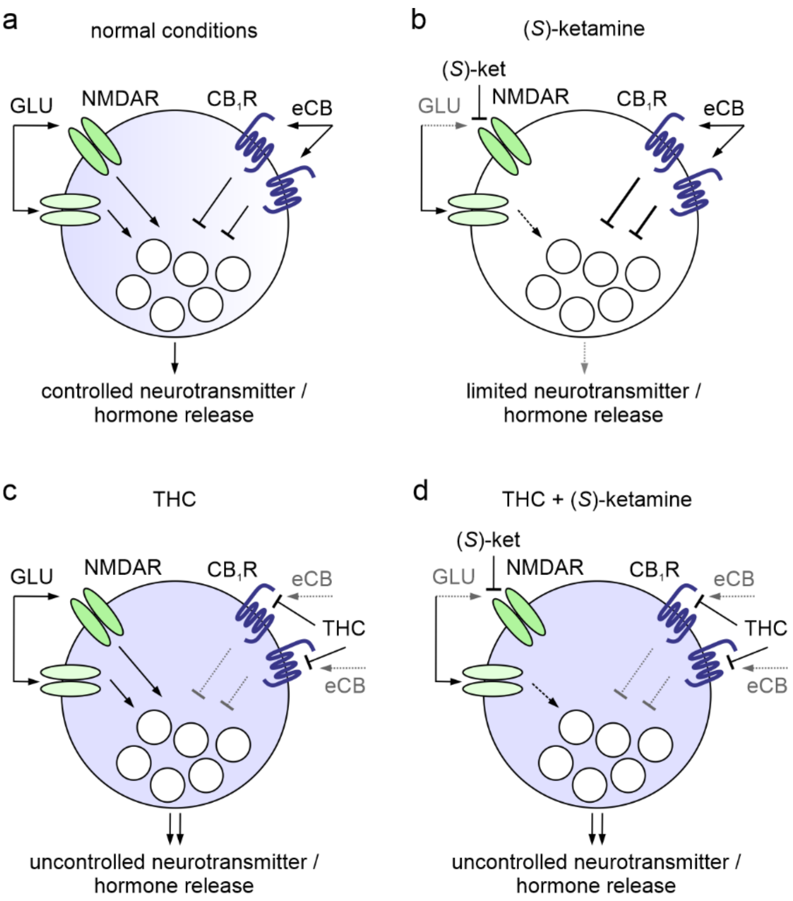


**Supporting Figure 7. Schema of drug interaction.** (**a**) Under physiological conditions, activation of NDMARs (*shades of green*) by glutamate (GLU) induces neurotransmitter and hormone release, while endocannabinoid (eCB) signaling through $CB_1$Rs reduces vesicular exocytosis for homeostatic control[1]. (**b**) As (*S*)-ketamine inhibits different NMDAR subtypes unevenly based on the (*S*)-ketamine dose provided[2], it could limit, but not completely prevent, the glutamate-induced release of neurotransmitters and hormones at both therapeutic and recreational doses. Thus, endocannabinoid effects could become dominant leading to a net reduction in neurotransmitter release. (**c**) In the presence of THC, a low affinity partial $CB_1$R agonist, the eCB-mediated control of cellular release becomes disrupted, leading to an increased net output of neurotransmitter and hormone availability[1]. (**d**) Upon THC exposure, (*S*)-ketamine is no longer able to diminish cellular release due to the lack of eCB-mediated control. *Abbreviations:* $CB_1$R, type 1 cannabinoid receptor; eCB, endocannabinoid; GLU, glutamate; NMDAR, N-methyl-D-aspartate (NMDA) receptor; (*S*)-ket, (*S*)-ketamine.

### Supporting Movies

**Supporting Movie S1 and S2.** Video recordings of catalepsy bar attempts of a successful 5 mg/kg THC (S1) and failed 10 mg/kg THC (S2) trial. Note that THC-treated mice at doses ≥ 5mg/kg exhibit a strong stereotypical startle response when placed on the bar, making reliable quantification complicated (*see also* Ref.[3]).

**Supporting Movie S3.** Video demonstrating the prolonged motor deficits (120 min post-THC) when (*S*)-ketamine is given 30 min after THC exposure. Note the slow and uncoordinated (hind leg) movement while traversing the bedding, a feature not observed when given THC or (*S*)-ketamine alone.

### Supporting References


1. Hillard, C. J., Beatka, M. & Sarvaideo, J. Endocannabinoid Signaling and the Hypothalamic-Pituitary-Adrenal Axis. *Compr Physiol* **7**, 1–15 (2016).
2. Zorumski, C. F., Izumi, Y. & Mennerick, S. Ketamine: NMDA Receptors and Beyond. *J Neurosci* **36**, 11158–11164 (2016).
3. Metna-Laurent, M., Mondésir, M., Grel, A., Vallée, M. & Piazza, P.-V. Cannabinoid-Induced Tetrad in Mice. *Current Protocols in Neuroscience* **80**, 9.59.1-9.59.10 (2017).
